# Exploration of the EEG response to periodic thermal and vibrotactile stimuli

**DOI:** 10.1101/2024.01.15.575576

**Authors:** Arthur S. Courtin, André Mouraux

**Affiliations:** Center for Functionally Integrative Neuroscience, Aarhus University, Aarhus (Denmark); Institute of NeuroScience, Université catholique de Louvain, Brussels (Belgium)

## Abstract

Under certain conditions, a stimulus applied at a given frequency will lead to a periodic variation of neural activity at the same frequency. Taking advantage of this periodicity, it is possible to tag this response in the EEG frequency spectrum. Frequency tagging of sustained periodic noxious heat stimuli led to the recording of phase-locked and non-phase-locked responses whose functional significance remains unclear.

This study aimed at assessing whether such responses can also be recorded during the repetitive presentation of brief innocuous cold, noxious heat and vibrotactile stimuli. Comparison between the responses obtained with different stimulation modalities should inform us on the nature of the neural processes underlying these responses (modality aspecific, somatosensory, thermosensory, nociceptive). Comparison between upper and lower limb stimulation should inform us on the somatotopic organization of these responses and, therefore, on their potential sources.

Based on our results, on one hand, trains of brief innocuous cold, noxious heat and vibrations can elicit phase-locked and non-phase-locked responses which appear highly similar to those evoked by sustained periodic noxious heat stimuli when frequency tagged. On the other hand, when analysed in the time domain or using time-frequency decomposition, these responses appeared highly similar to those that can be recorded following isolated brief noxious heat or tactile stimuli. These responses consisted in phase-locked activity corresponding to the vertex potential, thought to reflect modality non-specific attentional processes, and in an alpha-to-beta ERD originating in the S1/M1 area contralateral to the stimulated hand, probably reflecting non-specific somatosensory activity.

## 1. Background

Under certain conditions, a stimulus applied at a given frequency will lead to a periodic modulation of neural activity at this frequency. This can in turn be expected to translate to a periodic response in the EEG recorded during stimulus presentation. Taking advantage of this known periodicity, the EEG response can be “tagged” in the frequency spectrum of the EEG recording, as it will be concentrated at the frequency of stimulation and its harmonics. This “frequency tagging” approach has the advantage of suppressing the need for visual identification and extraction of event related potentials components, making response identification more objective and allowing to probe subtle cortical processes that would not lead to strong EEG responses [1, 2].

Attempts to use frequency tagging of nociceptive stimuli delivered at relatively high frequencies (5-6 Hz), like those used for other sensory modalities, yielded modest signal to noise ratios, probably because of skin nociceptors proneness to habituation/sensitization and activity dependent slowing [3–6]. The introduction of a new slow and sustained periodic noxious heat stimulus (5 s sinusoidal skin temperature elevation and cooling, from baseline skin temperature to 50°C) by Colon, Liberati [6] allowed to circumvent these problems.

The periodic activity they identified with these sustained painful heat stimuli consisted in central symmetrical phase locked activity, distributed around the vertex, and alpha (8 - 12 Hz) and beta band (12 – 30 Hz) oscillatory activity modulation, predominantly distributed in the centro-parietal region contralateral to the stimulated limb [6]. They also reported a periodic modulation of activity in the theta band with a less clearly defined topography. Using slightly different noxious heat stimuli, Mulders, de Bodt [7] observed EEG responses similar to those reported by Colon, Liberati [6].

The functional significance of these EEG responses remains unclear. Colon, Liberati [6] showed that the responses they recorded relied on the activation of unmyelinated primary afferents. However, this specificity at the periphery does not imply specificity of the brain responses. Given its topography, the phase-locked activity was hypothesized to correspond to the vertex potential that can be recorded in response to brief thermonociceptive stimuli but also to stimuli of other sensory modalities (touch, auditory, visual…) and which is thought to reflect mainly attentional processes [8, 9]. The periodic alpha and beta band modulation, on the other hand, could correspond to alpha-to-beta event related desynchronization (ERD) that can be recorded over the primary somatosensory cortex after brief noxious or tactile stimuli and is thought to reflect the activation of that brain area [10, 11].

Interestingly when frequency tagging a cold stimulus that was designed to mirror the noxious heat stimulus proposed by Colon, Liberati [6] (0.2 Hz sine wave between 32°C and 17°C), Mulders, de Bodt [7] obtained only weak and inconsistent EEG responses. It is however unclear whether this discrepancy between cold and heat evoked responses reflects specificity of these frequency tagged responses for noxious heat/pain processing or the inability of that cold stimulus to strongly and periodically activate cold sensitive afferents.

This study aimed at better understanding the nature of the periodic non-phase-locked responses identified by frequency tagging slow noxious heat stimuli. As alpha-to-beta ERD originating in S1 were previously reported following both brief tactile and noxious stimuli, we investigated the responses to different somatosensory modalities (innocuous cold, noxious heat, and vibration), to be able to disentangle processes related to somatosensation, thermosensation and nociception. If S1 is indeed involved in their generation, responses elicited by upper or lower limb stimulation should have distinct topographies, reflecting the somatotopic organization of that brain region. To test this hypothesis, we stimulated both the volar forearm and the foot dorsum. In order to rule out insufficient afferent pathway activation, stimuli used to record regular event related potentials (ERP) were used [12, 13], presented in trains delivered at a slow rate (to mimic the structure of the stimulus used by Colon, Liberati [6]).

## 2. Methods

### 2.1. Participants

Twenty healthy young volunteers (5 males & 15 females; age as mean ± standard deviation: 23.5 ± 2.5) each took part to two experimental sessions. One of these participants was excluded due to excessive environmental noise in the EEG recordings. Volunteers had to satisfy the following inclusion criteria: not having medical history of neurological, psychiatric, or metabolic diseases; not having a drug consumption habit (recreative drug or medicine use, alcohol abuse; anticonception drugs not included); not suffering of chronic pain; not having forearm/foot dorsum skin lesions or dermatological conditions; being aged between 18 and 65 years old.

All subjects gave their written informed consent prior to the beginning of the experiment. The study was approved by the local ethical committee and complied with the latest Helsinki Declaration (October 2013).

### 2.2. Thermal stimulations

Thermal stimuli were delivered using a Peltier effect contact thermode (TCSII, QST.Lab, Strasbourg, France). The stimulation surface of the TCS probe (model T03) is a disk (ø= 30 mm) divided in five independently controlled zones each set with three micro-Peltier elements (dimension of one element: 7.7 mm²). The stimulator enables cooling or heating the skin between 0°C and 60°C at a very high rate, with ramps of up to ±300°C/s. The delivered temperature is controlled at a rate of 1000 Hz by thermocouples located at the centre of each zone (resolution: 0.1°C).

Two different thermal stimulations were delivered to the participants (see figure 1): a train of cooling and a train of heating pulses, both delivered at 0.25 Hz and lasting a total of 60 s (15 periods). In both cases, all 5 zones were active for each pulse (total stimulation surface area: 116 mm^2^).

**Figure 1.**
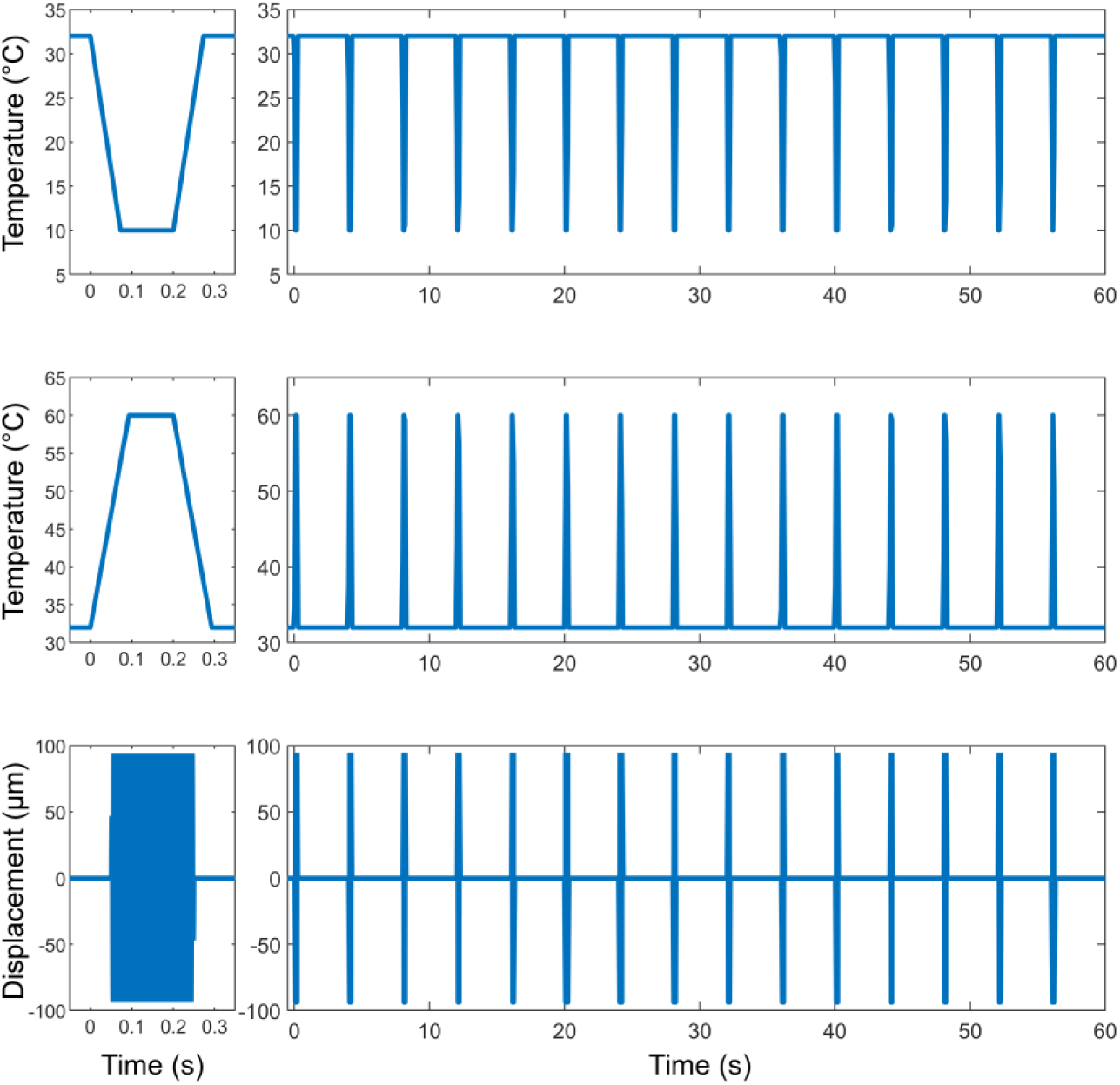
Stimulation profiles. This graph illustrates the individual impulses (left column) and complete stimulation profiles over time (right column) for the different modalities (first row: cold, second row: heat, third row: vibration). Temperatures are expressed in °C, vibration amplitude in µm and time in s.

**Figure 2.**
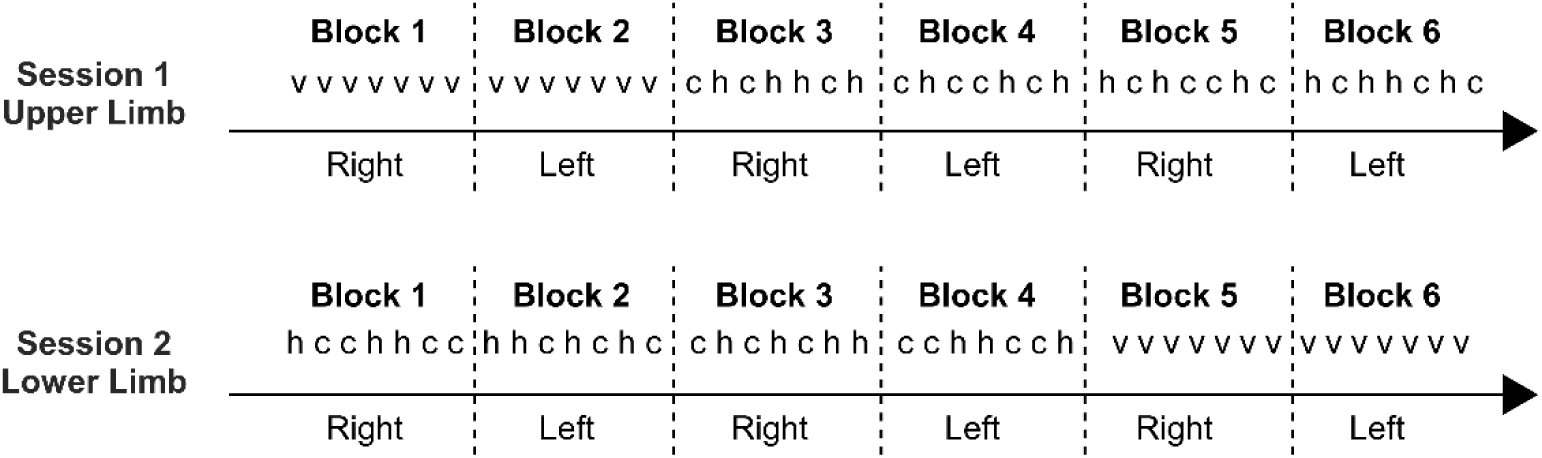
Example of block and stimuli order for one participant. v: vibrotactile stimulus, c: cold stimulus, h: hot stimulus

For the cooling pulses, the 5 zones dropped from 32°C to 10°C in ∼74 ms and remained at 10°C until 200 ms had elapsed since the beginning of the cooling. Then, the thermode heated back to 32°C in ∼74 ms and stayed at this temperature for the remaining of the 4 s period.

For the heating pulses, the active zones raised from 32°C to 60°C in ∼94 ms and remained at 60°C until 200 ms had elapsed since the beginning of the heating. Then, the thermode cooled back to 32°C ∼94 ms and stayed at this temperature for the remaining of the 4 s period.

### 2.3. Vibrotactile stimulations

Vibrotactile stimulations were delivered using the VTS 1 (Arsalis, Belgium), a 20 mm round-tipped piezo-electric actuator. Fifteen pulses of vibration (200 Hz), each lasting 200 ms, were delivered at a rate of 0.25 Hz, the stimulator being inactive the rest of the time (see figure 1). The surface of the stimulator was oscillating within a range of ±94 µm.

### 2.4. EEG recording

The EEG was recorded using a Biosemi Active-Two system (Biosemi) with 64 Ag-AgCl electrodes placed on the scalp according to the international 10/10 system. The signal was referenced to the CMS (Common Mode Sense) electrode and digitized at a 1024 Hz sampling rate. The EOG was recorded with 2 additional electrodes: one pasted to one of the zygomatic processes and the other between the eyebrows.

### 2.5. Procedure

Each participant was invited for 2 sessions, each lasting approximatively 2 hours. In one of them, the participant received stimulations on the volar forearms and, in the other, on the foot *dorsa*. Both sessions took place in the same dimly lit room. The participant was seated in a comfortable chair. The volar forearms were preferred to the hand dorsum, for upper limb stimulation, as, due to differences in the underlying tissues, they allowed a better contact between the skin and thermode.

After setting up the EEG cap, the participant was familiarized with white noise, delivered through ER-2 Insert Earphone system (Etymotic Research, the USA). The volume was set to the highest volume comfortable for the participant. For each participant and session, a vibrotactile stimuli was launched without the stimulator being in contact with the participant’s skin, to make sure that the white noise volume was enough to cover the noise generated by the stimulator. This was always the case. Then, one stimulus of each modality was presented on the hand dorsum of the participant, to familiarize them with the different stimulations they would receive.

During EEG acquisition, the participant was first presented with either vibrotactile or thermal stimulations. The stimulations were split in blocks of 7 stimuli. One block of vibrotactile stimuli was delivered on each side. Two blocks of thermal stimuli were delivered to each side. Hot and cold stimuli alternated in a pseudo random order within the block, guaranteeing that each side received a total of 7 vibrotactile, 7 cold and 7 hot stimuli (see figure 1 for an example).

The limbs stimulated in the first session, the side first stimulated, and whether the experiment started with vibrotactile or thermal stimuli were factors balanced between participants.

### 2.6. EEG Data Pre-processing

Processing and analysis of the EEG signal was conducted offline in MATLAB (The MathWorks) using the opensource toolbox Letswave 7 (https://www.letswave.org/). Channels Iz, P9 and P10 were excluded from the datasets as, due to their location, they were likely contaminated by muscular artefacts. The sampling rate of the data was decreased to 512 Hz. A FFT multi-notch filter was used to remove band line noise (notch frequency: 50 Hz, notch width: 2, slope width: 2, number of harmonics: 5). A 0.1-40 Hz band-pass Butterworth filter (order 4) was applied to the continuous EEG. The recordings were then segmented in epochs covering the 60 s of the stimulation. All electrodes were averaged-referenced by subtracting, for each time point, the mean across all channels from each channel. A validated method [14] based on an Independent Component Analysis (FastICA algorithm) was used to remove artefacts due to ocular activity. When ICs clearly reflected electrode noise, they were also removed. The number of excluded ICs *per* participant ranged from 3 to 12 out of 60 and was on average 8.9 (SD=2.1). Lateral scalp electrodes recorded during right side stimulations were flipped such that, regardless of whether the left or right hemi-body was stimulated, even-labelled electrode labels were contralateral to the stimulated side. Epochs containing amplitude changes exceeding 150 µV were rejected, as they were probably contaminated by artefacts. After this step, 13.55±0.83 out of 14 trials (mean ± SD) remained for cold stimulation of the foot, 13.70±0.92 for heat stimulation of the foot, 13.45±1.27 for vibrotactile stimulation of the foot, 13.95±0.22 remained for cold stimulation of the forearm, 13.80±0.41 remained for cold stimulation of the forearm, and 13.75±0.79 remained for cold stimulation of the forearm.

### 2.7. Frequency domain analysis

The data obtained at this stage constitute a first dataset, used to assess the existence of a periodic response in the phase locked EEG signal. Additional datasets were created to evaluate the effect of the periodic stimulation on the oscillatory activity present in the EEG. To do so, the signal was FFT multi-notch filtered to remove the components at the frequency of stimulation (FoS) and its harmonics (notch frequency: 0.25, notch width: 0.01, slope width: 0, number of harmonics: 1024). Several datasets were then created by bandpass filtering (order 8 Butterworth filter) the data to extract activity in the theta (4-8 Hz), alpha (8-12 Hz), and beta frequency bands (12-30 Hz). A Hilbert transform was then used to estimate the envelope of these oscillations as a function of time.

All datasets (original EEG signals to disclose phase-locked responses and Hilbert-transformed filtered signals to disclose non-phase-locked responses) were then averaged across epochs *per* stimulus, *per* limb and *per* participant. The obtained averages were segmented to start 3 s after the onset of the stimulus and to last until 1 s before the end of the stimulus, in order to remove the extremities of the signal (subject to Hilbert transform border effects). They were then transformed in the frequency domain using a Fast Fourier transform (FFT), yielding an amplitude spectrum (µV) ranging from 0 to 256 Hz. To obtain valid estimates of the magnitude of the EEG responses at each frequency, the contribution of noise was partially removed by subtracting, at each frequency bin and electrode, the average amplitude of the signal measured at neighbouring frequencies (*i.e.,* ±1 to 10 bins).

The resulting spectra were then segmented in pieces of 0.25 Hz centred on the frequency of stimulation (FoS) and its harmonics and summed to reassemble the periodic response (split between the FoS and its harmonics, as unitary response shape is not a perfect sinus; a total of 1024 segments were summed). This procedure resulted in vectors (13 bins/vector) for each channel/condition/participant combination, the central bin corresponding to the amplitude of the periodic response at the FoS and the other bins to random noise. Summing amplitude at harmonics to reconstruct the response amplitude is a common procedure in the frequency tagging literature [2].

To assess whether a response was present at the frequency of interest, multi-sensor cluster-based permutation one sample t-tests were used (using a right sided Wilcoxon sign rank test against 0, as the data was not normally distributed; number of permutations: 2000; alpha threshold: 0.05/6; cluster threshold: 0.05/6).

### 2.8. Time domain analysis

The dataset obtained at the end of the EEG data pre-processing were also segmented in epochs of 4 s, starting 1s before and finishing 3 s after the onset of the period/pulse. Epochs corresponding to the first period of each stimulus were discarded, as strong habituation of the ERP is expected between the first and the second presentation of a given pulse. The remaining epochs were averaged *per* participants. To account for inter-individual variability in the shape of the waveform, the group level representation of the-phase locked periodic response was build using 500 average waveforms from which 4 randomly chosen participants had been excluded.

### 2.9. Time frequency analysis

The datasets obtained at the end of the EEG data pre-processing were also analysed using time frequency transforms. First each epoch was cropped to range from 2.5 s to 59.5 s after the onset of the stimulation, in order to accommodate for the Hanning window bleeding at the extremities of the signal. Then, a windowed FFT was used to compute time frequency maps of the signal (Hanning width: 0.25 s, sliding step 0.064 s, frequency range: 4 to 30 Hz, frequency resolution: 1 Hz). The time frequency maps were then segmented in 4 s epochs and averaged across epochs.

All TF maps were baseline corrected by subtracting the average of the first second of the signal (corresponding to the second before period onset) for each frequency separately. The TF maps were then statistically tested against 0 using a bilateral Wilcoxon sign rank test (number of permutations:2000; alpha threshold: 0.05/6; cluster threshold: 0.05/6) to assess the presence of consistent modulations of the frequency content of the signal across participants.

## 3. Results

### 3.1. Frequency domain analysis

As can be seen in figure 3 all conditions led to significant time-locked and phase-locked periodic activity as well as a periodic modulation of the activity in theta, alpha and beta frequency bands (even though it was much weaker for lower limb thermal stimulations).

**Figure 3.**
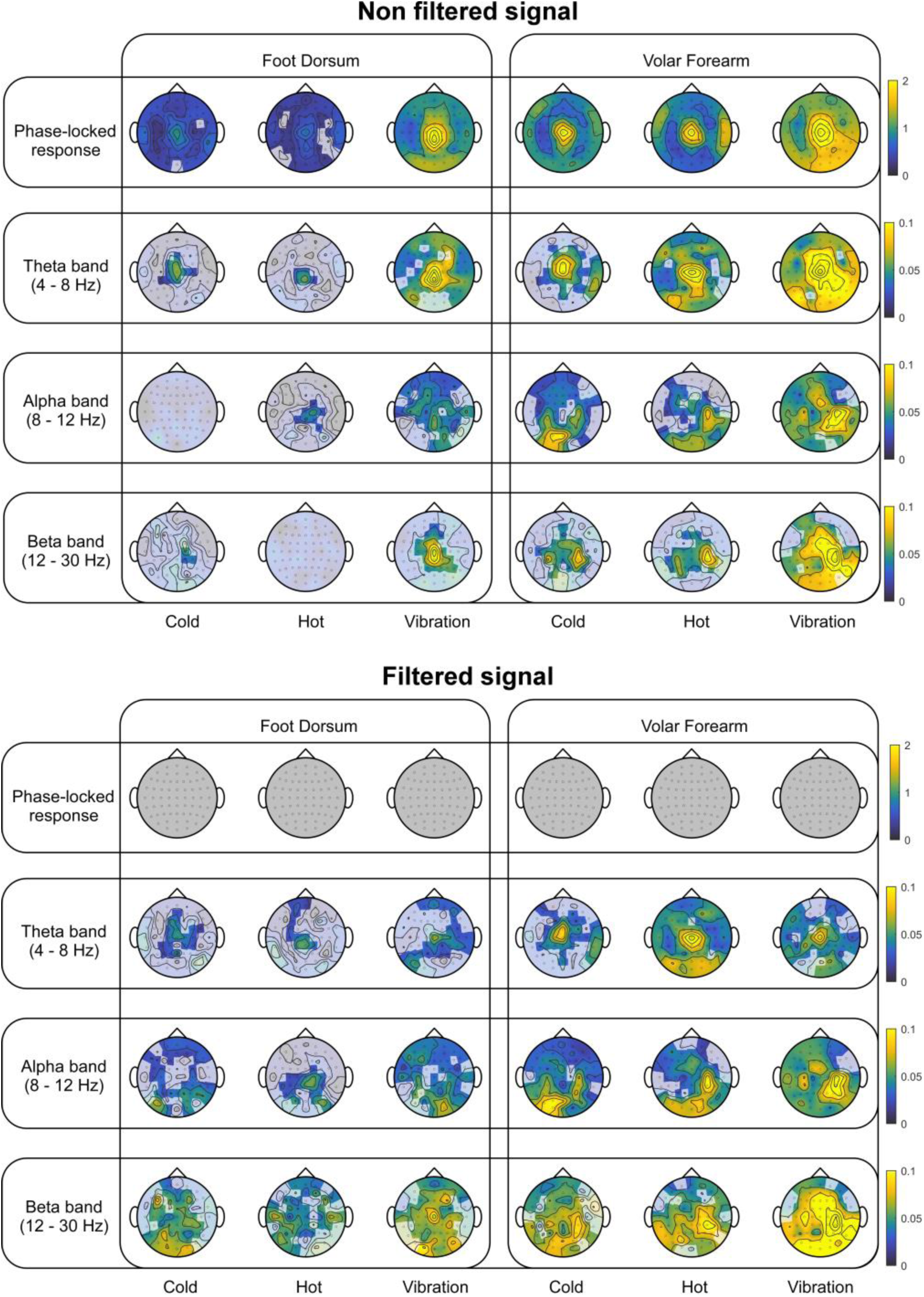
Group level median topographies of the periodic activity. For each electrode, the group level median value was plotted. A grey translucent shade was superimposed on electrodes that did not display significant periodic activity. This was tested by a right sided multi-sensor cluster-based permutation one sample Wilcoxon sign rank test.

The phase locked responses seemed mostly centred at the vertex with a possible lateralized additional component for vibrotactile stimulation of the forearm. As expected, no significant periodic activity could be detected in the signal filtered to remove activity at 0.25 Hz and its harmonics.

Overall, the theta band modulation matched the phase-locked activity scalp distribution. Filtering the signal to remove activity at 0.25 Hz and its harmonics led to a strong reduction of theta band modulations, indicating a significant contribution of the phase-locked responses to the activity in that frequency band. The remaining theta band modulations were also more or less centred around the vertex.

For forearm stimulation, activity in the alpha band seemed to centre around the centro-parietal region contralateral to the stimulated limb (C4, CP4) and possibly around the occipital region (especially for thermal stimuli) and the centro-parietal region ipsilateral to the stimulated hand (C4, CP4). For foot stimulations, the alpha band modulations were weak and the topographical pattern was less well defined, with possible clusters in the occipital and central region. Filtering the signal had little effect on activity in the alpha band.

Beta band modulations elicited by forearm stimulations seemed to distribute mainly over the centro-parietal region contralateral to the stimulated hand (C4, CP4) and in the occipital region. Similar to alpha band modulation, the topographic pattern was less clearly defined following foot stimulation, with possible foci in the occipital region and around the vertex.

### 3.1. Time domain analysis

As expected, all stimulation modalities led to clear group level periodic ERPs (see Figure 4, blue traces). These responses were in the Aβ conduction velocity range for vibrotactile stimuli (compatible with low threshold mechanoreceptor activation), in the Aδ range for cold stimuli (compatible with cold thermoreceptor activation) and both in the Aδ and C conduction velocity range for heat (compatible with the activation of both myelinated and unmyelinated heat sensitive nociceptors). Such bimodal responses to noxious heat stimuli were already observed in a previous study using the same stimulator and similar unitary stimuli [15].

**Figure 4.**
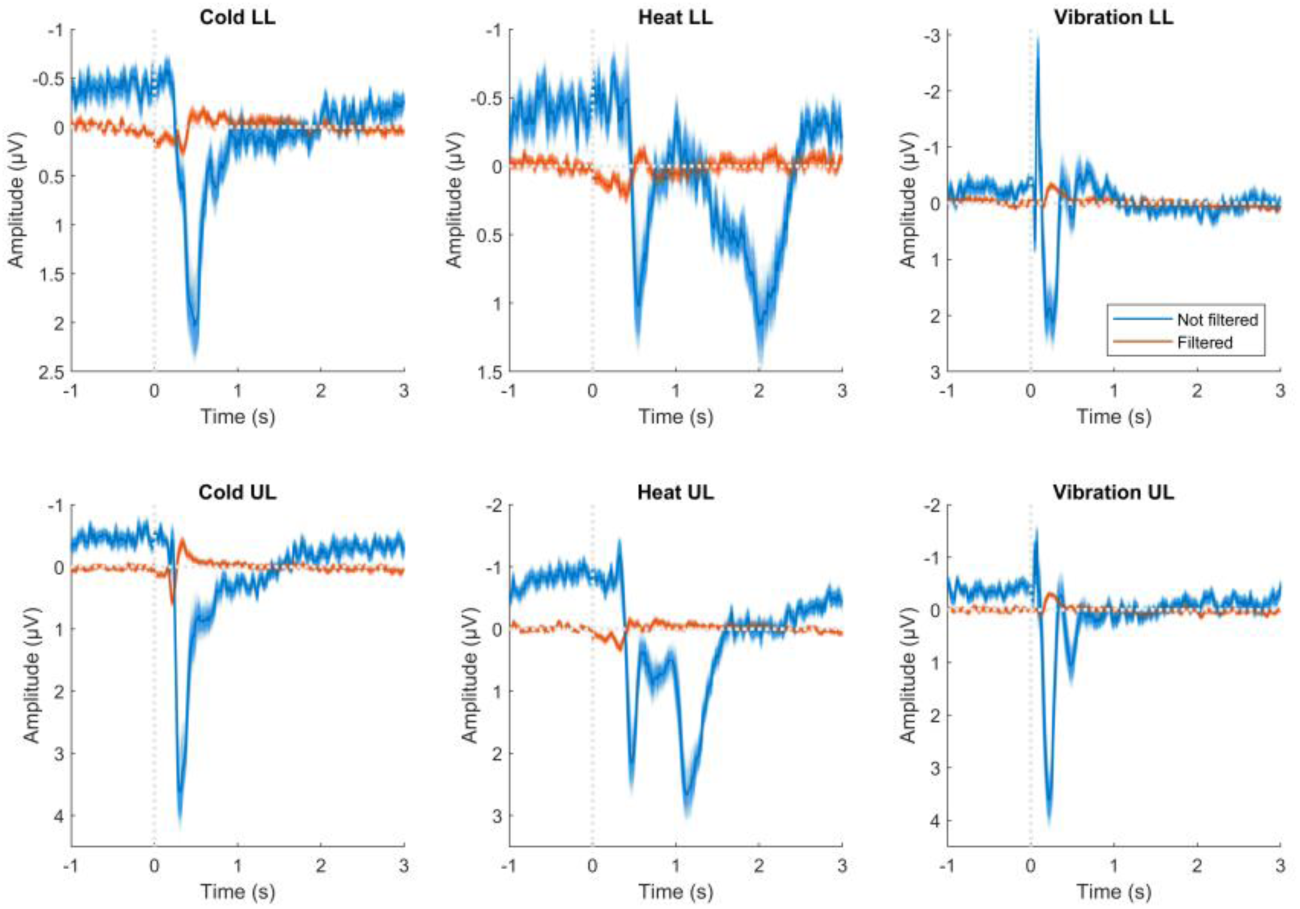
Group level representation of the phase locked periodic response at the vertex (Cz electrode) The darker lines represent group level averages. The shadow around them was constructed using 500 iterations of a leave 4 out procedure, i.e., for each iteration the data of 4 randomly chosen participants were left out of the dataset before the across subject average was computed. This procedure allows to represent the group level variability. The blue lines represent the regular dataset, used to assess phase-locked responses, and the orange line represents the dataset to which a multi-notch FFT filter was applied to remove components at 0.25 Hz (frequency of stimulation) and its harmonics. As can be seen from the orange traces, this filter did not completely remove the periodic evoked potentials triggered by the stimulation, probably because of temporal jitter of the individual responses.

Based on the group level averaged waveforms of the filtered datasets (see figure 4, orange traces), it seems that the multi-notch filter used to remove phase locked activity at the frequency of stimulation was not able to remove it entirely.

### 3.2. Time frequency analysis

The time frequency maps (see figure 5) show an initial increase of power in the theta to beta range which seem to correspond to the phase-locked response, as it is consistently present only in the non-filtered signal.

**Figure 5.**
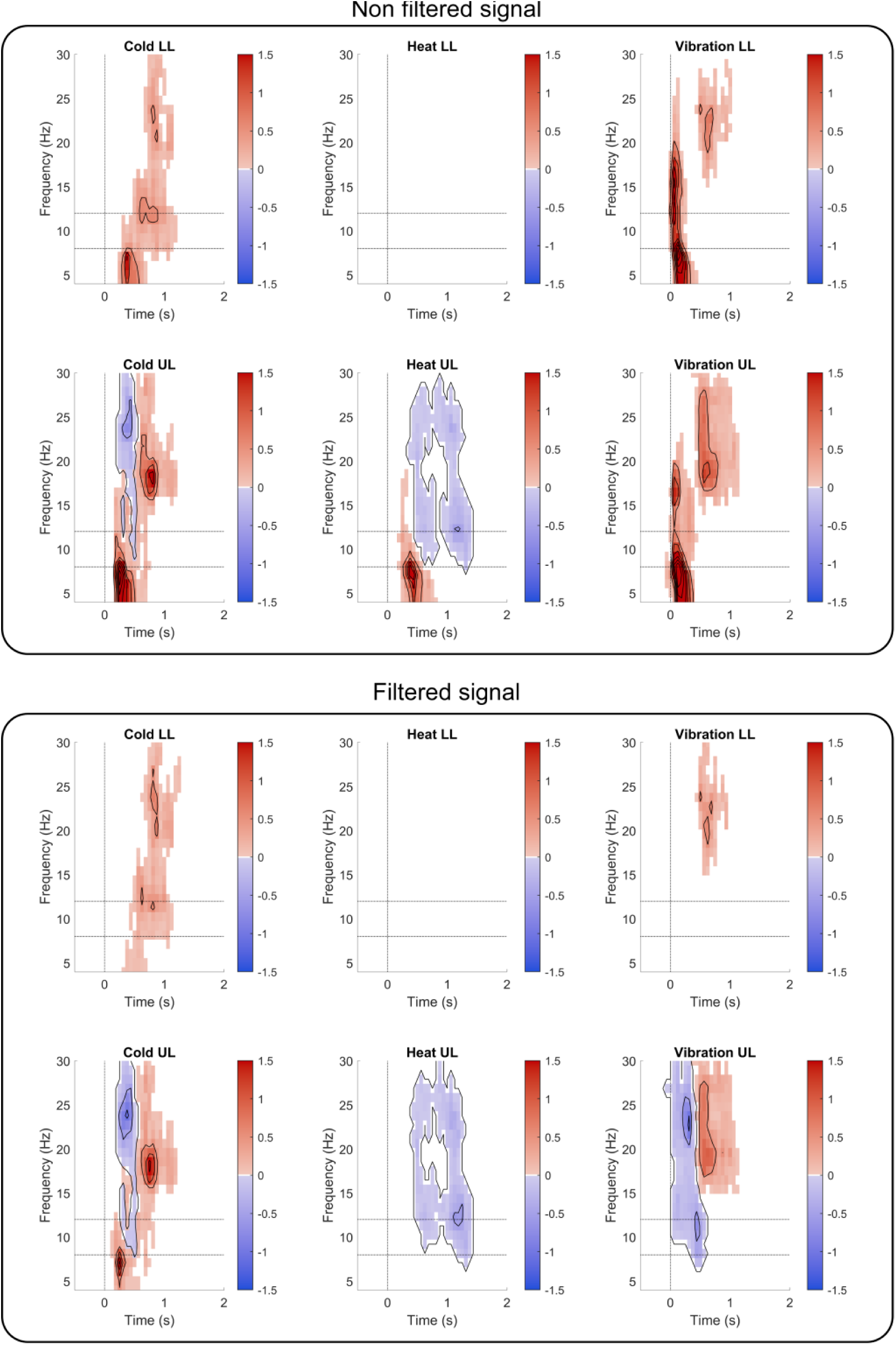
Time frequency decomposition of the periodic response. The plotted values summarise the average across electrodes of significant increases and decreases in the EEG activity across time and frequency. To construct this figure, the z-value approximation of the Wilcoxon sign rank test statistic that survived the multi-sensor cluster-based permutation correction were averaged across electrodes. The upper part of figure corresponds to dataset with the entirety of the signal whereas most of the phase-locked activity was removed prior to the computation of the time frequency maps represented in the lower part.

Cold stimuli led to an increase of theta band activity that survived the filter, in the same time-range as the phase-locked responses described in the previous paragraph.

Upper limb stimuli, led to decreases of power in the alpha band, appearing at about the same time as the phase-locked responses. Interestingly, for noxious heat, these decreases reflected the biphasic nature of the phase-locked responses. These alpha band modulations seemed to mainly take place over the centro-parietal region contra-lateral to the stimulated limb (see figure 6). In the case of innocuous cold stimuli, this was also accompanied by an increase of alpha band activity over the occipital region (especially in the side ipsilateral to the stimulated limb) and, for lower limb stimulations, over the ipsi and contralateral centro-parietal regions.

**Figure 6.**
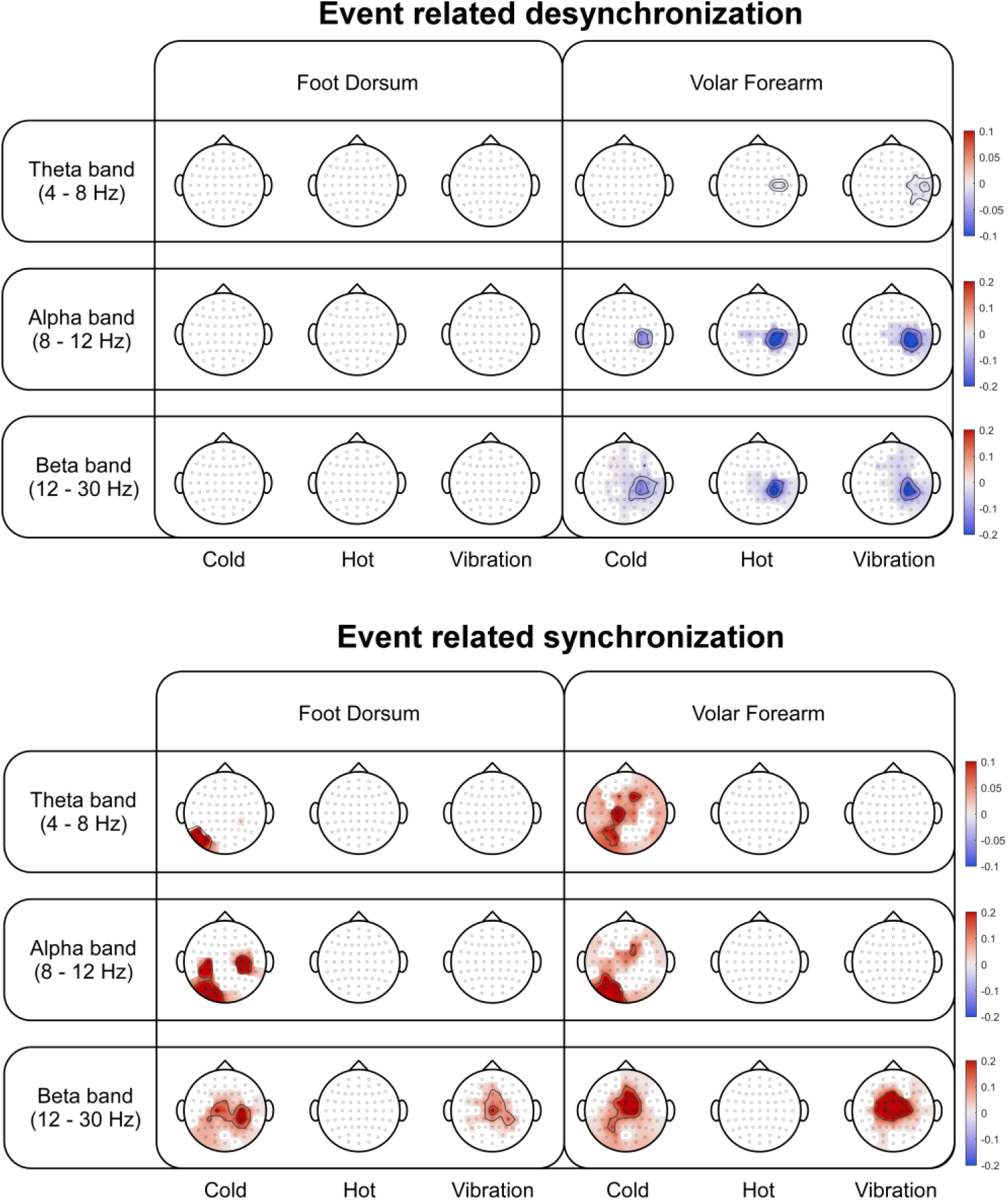
Topographies of the event related synchronization and desynchronization revealed by the time frequency analysis. To construct these topographies, the z-value approximations of the Wilcoxon sign rank test statistic that survived the multi-sensor cluster-based permutation correction were averaged across time and frequency (within the limits of each band). This was done separately for significant increases and decreases of activity.

Concomitantly to the alpha band activity decrease, reduction of beta band activity was also observed for upper limb stimulations, with a similar topography (even though less focused). This was followed by an increase in beta band activity for cold and vibrotactile stimulations, both for upper and lower limb stimulations, which seemed centrally distributed.

## 4. Discussion

### 4.1. Interpretation of the evoked responses

#### 4.1.1. Phase locked response

The frequency tagged phase-locked responses consisted primarily of activity centred at the vertex for all limbs and modalities. This is consistent with the type of responses recorded by Colon, Liberati [6] and Mulders, de Bodt [7]. No significant phase-locked periodic activity survived the statistical test after filtering. In the time frequency analysis, filtering the signal led to the disappearance of an early increase of activity spanning the theta to beta range, whose latency was compatible with the event related potentials that can be observed in the time domain. The topography and behaviour of these responses is highly compatible with the idea that the frequency tagged activity reflects the classical ERPs that can be recorded after brief thermal and vibrotactile stimuli and in particular the vertex potential. This ERP component can be recorded after the delivery of sensory stimuli pertaining to a large variety of modalities (somatosensory, visual, auditory…) and is thought to reflect mainly modality aspecific attentional processes [8, 9].

Interestingly, vibrotactile stimulations of the upper limb also seemed to elicit lateralized periodic activity. The distribution of this lateral activity is similar to that of early components of the non-nociceptive ERPs and that of frequency tagged activity elicited by vibrotactile stimuli modulated at a relatively high frequency [4]. Both are thought to reflect primary somatosensory cortex activity. The fact that this lateralized component seems to be absent from recordings obtained during foot dorsum stimulations could be explained by the position of S1 foot representation which is much closer to the midline than the representation of the forearm, what could lead responses evoked by its activation to blend in the topography of the vertex potential.

#### 4.1.2. Theta band modulation

Frequency tagged periodic theta band modulations were strongly reduced by filtering the signal to remove phase-locked activity at the frequency of stimulation. Their topographies seemed to match those of phase-locked responses, even after filtering. As the filter was not able to perfectly remove the phase-locked activity, probably because the periodic response was not perfectly stable from cycle to cycle (temporal jitter, habituation/sensitization), it is possible that these remaining theta band modulations correspond to the part of the phase-locked response that was not filtered out (see figure 3).

Interestingly, even though both studies reported a significant modulation of theta activity, Colon, Liberati [6] and Mulders, de Bodt [7] did not report a clearly defined topography. Given that they used a very slow stimulus, it is likely that the periodic responses they recorded were less prone to jitter and that the activity corresponding to jitter would have had a lower frequency content, mostly below the theta band.

#### 4.1.3. Alpha and beta band event-related desynchronization

For upper limb stimulations, all three modalities led to an event related desynchronization (ERD) in the alpha and beta bands, clearly localized over the contralateral centroparietal region. This ERD most certainly explains the periodic activity recorded in that region using frequency tagging, in this study as well as in those of Mulders, de Bodt [7] and Colon, Liberati [6]. Given the similarity between modalities and given its localization, it seems likely that this ERD originates in the primary somatosensory (or motor) cortex contralateral to the stimulated forearm and reflects general somaesthetic processes.

Similar ERD could not be reliably recorded following foot dorsum stimulation, neither in the contra-lateral centro-parietal region nor anywhere else. This observation seems to further confirm an S1/M1 origin for the ERD evoked by upper limb stimulations. Importantly, it does not seem likely that the absence of ERD in the contralateral centro-parietal region following foot stimulation comes from a bad signal-to-noise ratio for responses elicited after lower limb stimulation. Indeed, whereas the phase-locked responses evoked by thermal stimulation of this extremity were weak, responses to vibrotactile stimulation of the foot appeared stronger than responses to thermal stimulation of the forearm.

Such somatosensory ERDs have been recorded in response to brief noxious, tactile and tactile-like electrical stimuli [11, 16–19]. This response has been shown to correlate with stimulus intensity and with task relevance of the somatosensory modality. Following the largely accepted view that alpha desynchronization reflects cortical disinhibition, this ERD could correspond to activation of S1/M1 by these stimuli [20].

Interestingly, as far as we know, cold-evoked somatosensory alpha-to-beta ERD had not been reported before. The existence of this ERD is compatible with the long held view that cold perception involves S1 activation [21].

#### 4.1.4. Occipital alpha modulation

Along the alpha-to-beta ERD over the sensory-motor cortex, phasic painful stimuli usually evoke a similar and more or less concomitant ERD over the occipital regions [16, 19]. Such a response has been associated with the allocation of attentional resources to task-related processes [19]. This could also be interpreted as a spontaneous increase of visual activity in response to painful stimuli, aimed at locating the stimulus source in order to react to the threat it causes.

Such a response was not observed in our data and, instead, cold stimuli led to an occipital alpha band ERS. Interestingly, an alpha ERS in the occipital region has been reported during tonic cold pain stimulations, which was interpreted as reflecting the attention being dragged away from the visual modality towards pain [22].

As the repetitive stimuli used in this experiment were highly predictable, it is possible that unitary stimuli (cold/heat/vibration pulses) did not consistently lead to attention reorientation or, at least, that this response quickly habituated over the 15 cycles of a full stimulus. Additionally, as participants were instructed to fix their gaze, it is also possible that the occipital alpha ERS that was observed following cold stimuli reflected active inhibition, counteracting this reflex reallocation of attention to the visual modality.

Interpreting this occipital alpha ERS as a specific marker of cold processing, because only this modality led to a significant response in that region, seems problematic. Indeed, when looking at the frequency tagged responses, occipital alpha modulations seem present for all modalities. This difference between modalities in the time-frequency analysis more likely originates from the statistical procedure used to highlight its existence. Indeed, the minimal detectable response is directly related to the maximum response under the H0. As can be seen in figure 6, the alpha ERD is weaker for cold than for heat or vibration and this could explain why the alpha ERS survive correction only for the first modality.

#### 4.1.5. Beta band event-related synchronization

Cold and vibrotactile stimuli also evoked a later beta band synchronisation that was widespread and diffuse but was particularly pronounced around the vertex. This response could correspond to the late/ultra-late beta ERS recorded by Mouraux, Guérit [23] after brief laser stimuli. The significance of this ERS is unclear and it could be a kind of beta rebound such as that observed over sensorimotor areas after the end of a movement [24]. Its central topography could result from many different generators. Interestingly, it seemed to be similar for upper and lower limb stimulations and we can therefore rule out that this activity would originates in the primary somatosensory or motor cortices.

### 4.2. Advantages and limitations of the frequency tagging framework to study thermosensation and thermonociception

Based on our data, it seems likely that the frequency tagged responses to sustained repetitive nociceptive stimuli recorded by Colon, Liberati [6] and Mulders, de Bodt [7] can also be recorded using time frequency analysis of brief stimuli. Indeed, their phase-locked responses seem to correspond to the ERPs elicited by brief stimuli of the same modality and especially the vertex potential and the contralateral alpha and beta band modulations seem to correspond to the sensorimotor alpha-to-beta ERD that is observed after brief noxious/tactile stimuli. Therefore, frequency tagging allows us to look at the same responses in a different way but not to look at different responses.

Additionally, frequency tagging implies collapsing the recorded signal over time. In doing so, it leads to a loss of information as it summarizes one period (several seconds of recording in our case) into one topography, agglomerating different responses/processes together. This lower definition of the outcome would probably result in decreased sensitivity to modulations of the brain response, as it is more difficult to detect small changes in one specific component of the response if all components are diluted together.

On the other hands, frequency tagging also has the interesting property that the distribution of signal and noise in the frequency spectrum is deterministically known. This has the advantage of removing experimenters’ degrees of freedom in the identification and extraction of brain responses, making the data extraction and analysis more objective.

Additionally, this also has the advantage of leading to the possibility of identifying EEG responses to stimuli which do not lead to well defined ERP/ERS/ERD. For example, the very slow stimuli used by Colon, Liberati [6] allow to record brain responses to the preferential activation of slow-adapting unmyelinated polymodal nociceptors. Due to their slow adapting nature and due to significant heat activation threshold variability in this primary afferent subpopulation, recording regular ERPs in response to their activation is not possible.

Alternatively, one could want to use this type of sustained stimuli not to study the brain reaction to a specific primary afferent input but because this type of sustained slowly modulated stimuli are more ecologically relevant to study pain processing than pulses of noxious heat lasting a few milliseconds.

Finally, this deterministic repartition of noise and signal in the frequency spectrum also allows to statistically assess the presence of a significant brain response to a given stimulus at the individual level, presenting interesting possibilities of n-of-1 design, for example to assess the effectivity of clinical interventions [2, 25, 26].

## 5. Conclusion

Frequency tagging the EEG responses to trains of brief innocuous cold, noxious heat or vibration pulses lead to the same kind of responses across modalities. These responses were also very similar to those recorded by previous studies who used sustained slowly modulated noxious heat stimuli. The phase-locked responses to these stimuli appeared to mainly correspond to the vertex potential, thought to reflect modality non-specific attentional processes related to sensory perception. The alpha and beta periodic activity appeared to reflect an event related desynchronization, likely corresponding to the activation of S1 by these stimuli.

## Acknowledgement

ASC was a FRIA grantee of the Fund for Scientific Research – FNRS at the time of data collection and during part of the analysis and writing process.

